# Evaluating Cross-linking-driven integrative modeling in peptide-HLAII complexes prediction with insights for refining predictive accuracy

**DOI:** 10.1101/2024.09.03.609139

**Authors:** Andrea Di Ianni, Luca M. Barbero, Kyra Cowan, Federico Riccardi Sirtori

## Abstract

*In silico* prediction of peptide-HLAII (human leucocyte antigen class II) complexes has emerged as a crucial approach in bioinformatics for deciphering antigen presentation mechanisms. Several *in silico* tools have been developed to predict peptide binding to HLAII alleles, trying to deconvolute the intricate peptide-HLAII binding specificity. These approaches integrate bases from molecular modeling, machine learning, and bioinformatics to predict peptide-HLAII interactions. Initially, structure-based methods relying on molecular docking algorithms were widespread, utilizing structural data of HLAII molecules and peptides to infer plausible binding conformations. These methods often faced challenges in accuracy due to the dynamic nature of peptide-HLAII interactions. Besides, the high flexibility of peptide sidechains makes their placement into the HLA-binding site even more complex. In recent years, machine learning techniques have drawn attention to peptide-HLAII binding predictions. Supervised learning algorithms, such as support vector machines (SVMs), neural networks, and ensemble methods, have been considerably applied to discriminate patterns from large datasets of experimentally validated peptide-HLAII binding affinities (like Immune Epitope Data Base, IEDB) and more recently mass spectrometry-eluted ligands from MHC-associated peptide proteomics (MAPPs) assay. The role of experiment- assisted integrative modeling in aiding peptide-HLAII complexes prediction still needs to be clarified. In this work, we benchmarked the use of AlphaLink2 (AlphaFold2 + cross-links restraints) and compared it to AlphaFold2 Multimer in predicting correct peptide binding motifs. These results can pave the way to an integrated strategy for vaccine development and protein deimmunization or autoimmunity mitigation.

## INTRODUCTION

The human leucocyte antigen (HLA), or more generally major-histocompatibility complex (MHC) class II molecules, are key receptors implicated in acquired immunity^1^. HLA class II molecules act against extracellular proteins, by cleaving them into small peptides to which they interact to activate the immune system^2,3^. Due to their role in managing infections caused by external pathogens and in the recognition^4^ of biologics by immune cells, understanding the HLA class II binding mechanisms^5^, and predicting the binding affinity to peptides and their structural binding motifs, has become a priority. The molecular mechanisms by which HLA class II recognizes peptides have been deeply investigated over the years^6^. The peptides bind to a groove located on the external regions of α1 and β1 chains of the HLA class II receptor, which is characterized by two α-helical walls and a β-sheet^7^. The binding peptides can have a length between 9 and 20 amino acids. However, the complex is mainly stabilized by the interactions of 9 amino acids in the peptide core, which are localized in four key pockets: P1, P4, P6 and P9. It has been reported that the interactions at the P1 region (including the P2 site) in HLA class II appear to have a dominant effect on the presentation of stable peptide-HLAII complexes and on the immunodominance of certain peptide epitopes^8^. The receptor key pockets are located between the α1 and β1 regions. The latter contains multiple polymorphisms, associated with different HLA class II DR alleles. HLA-DP and DQ molecules are polymorphic on both α and β chains of the peptide binding pocket^9,10^. The peptide flanking regions (PFRs), as well as the amino acids that do not interact with the HLA anchor points, are crucial to facilitate the interaction with T-cell receptors (TCRs) after migration of the peptide−HLA complex to the surface of antigen-presenting cells (APCs)^11^.

### Background on HLA Class II Molecules and their Importance in Immunity

HLA class II molecules bind peptides differently than their HLA class I counterparts. For example, peptides binding to HLA class II molecules could not be limited to both ends of the peptide-binding groove. The ends of HLA II molecules are open, explaining why these proteins can effectively engage 14-mer and longer peptides, whereas HLA class I are limited to relatively short (7–10 residue) peptides^12–14^. Then, even longer peptides, could be accommodated by protruding out of one or both ends of the HLA class II groove. This clarified why these peptides did not exhibit common anchor residues when aligned by their N termini. Moreover, conserved residues within the class II peptide-binding groove made hydrogen bonds with main chain atoms within the peptide. As a result, the peptide adopts a twisted conformation called polyproline II helix, where all peptide atoms are not busy with internal hydrogen bonds and are free for external interactions with the HLA binding site^7^. These interactions with peptide main chain atoms justified the capacity of each HLA II allele to bind different peptides. Instead, peptide-HLA II allele specificity is supported by peptide side chain interactions at anchor positions within the receptor groove. Identifying peptides that potentially trigger immune responses partially relies on the affinity of the interaction between the peptide and the HLA class II receptor^15^. Consequently, the higher the peptide affinity the greater its potential to trigger an immune response. Nevertheless, other components could affect the immunological recognition of these peptides, such as the stability of the interaction between the peptide and the HLA II molecule, the peptide−HLA complex complementary interaction with TCR loops, and the editing process made by additional chaperones like HLA-DM and DO^16–18^.

### Advancements in predictive tools for HLA-Peptide Binding

With this in mind, the area of immunoinformatics has developed a wide set of bioinformatics tools (mainly sequence-based tools)^19^ for predicting the affinity between a peptide and HLA class I and class II molecules^20,21^. These bioinformatics pipelines should detect if novel peptides derived from pathogen/exogenous proteins (e.g., viruses and bacteria, therapeutic proteins) can be used as vaccine candidates or can mitigate their immune response (in the drug discovery field)^21–23^. Initially, motif-based approaches were used to identify binding motifs in antigenic peptides, focusing on anchor points believed to stabilize the peptide-HLA interaction^24^. However, it was discovered that peptides without these specific motifs could still bind to HLA class II molecules due to secondary anchor positions and PFRs that influence their docking. With the advent of machine learning, numerous algorithms were developed^25^. These algorithms improved due to the increasing experimental data available for training. For MHC class II, various methods have been developed to replicate the MHC binding environment and peptide interaction using machine learning^26^. These methods aimed at HLA allele pan-specificity, covering all known HLA class II variants. Binding to MHC is the most selective step in antigen presentation, but other factors like antigen processing and peptide-MHC complex stability also play impactful roles. Algorithms based solely on MHC affinity can be over predictive, so additional filtering is necessary. Mass spectrometry (MS) peptidome data (such as MHC-associated peptide proteomics assay, MAPPs), which considers antigen processing and presentation, is promising for creating large-scale peptide libraries that characterize the potential ligands for individual MHC molecules^27–29^. However, there is an inequity in data coverage between binding assays (BA) and eluted ligands (EL), with the EL dataset being relatively smaller compared to the BA dataset. Besides, EL data is not as quantitative as BA data. To address this, prediction algorithms should incorporate both MHC binding and ligand elution data. NetMHCpan version 4.0 by Jurtz et al. achieved this by combining MHC binding data for 130 MHC class I alleles and MHC ligand elution data for 55 alleles^30^. They trained a single neural network with a novel architecture that predicts binding affinity and the likelihood of a peptide being an eluted ligand. Usually, most immunopeptidomics data have been generated by applying HLA-DR–specific antibodies during the HLA-bound peptides affinity purification step before the MS analysis. Eventually, pan–HLA class II antibodies or a mixture of HLA class II-specific antibodies can be applied to capture the residual class II molecules (HLA-DP and HLA-DQ) in the sample^31,32^. However, these antibodies have demonstrated poor affinity and specificity toward both DP and DQ, thereby having low peptide yield for these loci^33,34^. Another critical issue associated with the interpretation of MS analysis is that the data most often derive from multi-allelic (MA) cell lines, implying that each peptide can arise from the set of possible HLA molecules expressed in the given biological sample. Therefore, elucidation of this type of data requires a classification of the peptides into their most likely HLA restriction, a process known as motif deconvolution. Several methods have been proposed, such as MoDec^33^ and NNAlign_MA^34^. Recently, Nilsson et al. released the last update of NetMHCIIpan prediction method (NetMHCIIpan 4.3 version), with increased performance on all three HLA class II loci^35^. Firstly, they integrated large immunopeptidomics datasets covering all HLA class II locus specificity and refined the machine learning framework by including inverted peptide binders. Concerning molecular modeling prediction tools for protein-peptide interaction, the role of assisted integrative modeling in aiding peptide-HLAII complexes prediction still needs to be clarified. Cross-linking mass spectrometry (XL-MS) data can be used to drive integrative modeling, where cross-links information serves as distance restraints for the precise determination of binding interfaces^36^. In this work, we benchmarked AlphaLink2^37–39^ (AlphaFold2 + cross-links restraints) and compared it to AlphaFold2 Multimer^40,41^ in predicting correct peptide binding motifs. We then assessed the performance of each pipeline on the different HLA class II loci. We also investigated factors contributing to modeling accuracy such as peptide length and noncanonical peptide binding conformations. Our study presented a thorough assessment of AlphaLink2/AlphaFold2’s ability to predict peptide-HLA complexes, giving helpful insights for interpreting model accuracy and identifying potential key factors to consider when modeling these complexes. To our knowledge, this work represents the first application of XL-MS and integrative modeling in the immunopeptidomics field. In the next years, an integrated strategy of physics-based and sequence-based bioinformatics tools can be helpful to screen potential peptide mutants/candidates for vaccine development or assist the deimmunization process of exogenous proteins/peptides.

## MATERIAL AND METHODS

### Chemicals

All chemicals used in this study were of the highest purity available and were obtained from Sigma Aldrich (St. Louis, USA). Recombinant model CLIP/HLA-DRB1*01:01 complex (product code: TB- RD-300) was purchased from MBLI (Schaumburg, USA). Disuccinimidyl dibutyric urea (DSBU) was purchased from Thermo Fisher Scientific, Massachusetts, USA (Cat: A35459). 4-(4,6- dimethoxy-1,3,5-triazin-2-yl)-4-methyl-morpholinium chloride (DMTMM) was purchased from Sigma Aldrich (Cat: 74104). Nano-HPLC solvents were LC/MS grade (LiChrosolv®, Merck KGaA).

### Peptide-HLAII complex benchmark assembly

We assembled a set of high-resolution structures to benchmark AlphaFold2 and AlphaLink2 composed by a total of 55 structures (35 different peptide/HLA-DR, 10 peptide/HLA-DP and 10 peptide/HLA-DQ complexes). To obtain an initial list of peptide-HLA complexes, we downloaded the full set of peptide-HLA II structures dataset from UniProtKB (entries P01903, Q30154, P79483, P20036, P04440, P01906, P01920, P05538, P01909). From this initial list, the peptide-HLA complex dataset for AlphaFold2 and AlphaLink2 benchmarking was assembled using the following criteria:

(1) crystal complex structure resolution ≤ 3.0 Å, (2) non-redundant structures, (3) peptide with no post-translational modification (e.g., acetylation, deamidation, citrullination) or chemical modifications (because they cannot be predicted in current AlphaFold2/AlphaLink2 Colab Notebooks). Non-redundancy criteria for peptide sequence is sequence ID at least <90%. Different peptide lengths were included in the dataset. The AlphaFold2 benchmarking was performed with ColabFold version 1.5.5^37^, using scripts from Github. AlphaLink2 benchmarking was performed from the Colab Notebook on the same AlphaFold2 case studies, shown in Table S1.

### AlphaFold2 peptide-HLA modeling

Peptide-HLA complexes were predicted using the AlphaFold Multimer Colab Notebook (https://colab.research.google.com/github/deepmind/alphafold/blob/main/notebooks/AlphaFold.ipy nb) by entering the sequence of the HLA protein and peptide. AlphaFold was run in multimer mode with no PDB templates information, 3 number of recycles and 5 relaxation cycles. 5 models were generated for each complex, using mmseqs2_uniref_env as Multiple Sequence Alignment (MSA) mode.

### AlphaLink2 peptide-HLA modeling

Peptide-HLA complexes were predicted using the AlphaLink2 Colab Notebook (https://colab.research.google.com/github/Rappsilber-Laboratory/AlphaLink2/blob/main/notebooks/alphalink2.ipynb#scrollTo=RJUxaO7Ofw1L) by entering the sequence of the two proteins. AlphaLink2 was run using optimized model weights from experimental cross-linking data (10Å Cα-Cα cut-off). MSA was performed using MMseqs2 and without PDB templates. Max recycling iterations were set to 3 for model inference. 3 models were generated for each complex. Four different simulated cross-link datasets were used to perform predictions for each peptide-HLA case study (see details in Table S1). For experimentally-determined cross-links on a tested peptide-HLA complex, a .csv file with all identified interprotein cross-links was used as cross-links restraint file to perform the model prediction.

### Interface pLDDT calculation

To determine the interface predicted Local Distance Difference Test (I-pLDDT), we computed the average pLDDT value for all residues at the peptide-HLA interface. Interface residues were defined as any residue with a non-hydrogen atom within 8.0 Å of the binding partner, following Marcu et al. FlexPepDock peptide-protein CAPRI round study^42^.

### Ligand normalized model confidence

To determine the ligand normalized model confidence, a ligand normalization score (LNS) was calculated by using the following equation (Equation 1).

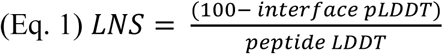

Equation 1. Ligand normalization score calculation.

This normalization score was used to derive the ligand normalized model confidence.

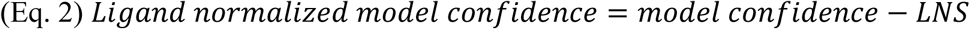

Equation 2. Ligand normalized model confidence calculation. Plots and statistical analysis

PyMOL^43^ (Schrodinger, Inc.) was used to generate peptide-HLA structural figures. The matplotlib package in Python was utilized to generate box plots, strip plots, and scatter plots. Pearson correlations and their corresponding p-values were calculated using the pearsonr function from scipy.stats package in Python. Shapiro-Wilk test for testing the normality of the data distribution was performed using Shapiro function from scipy.stats package in Python. Cullen and Frey graph was used to assess the distribution type of different model accuracy predictors using descdist function of fitdistrplus package in R. Wilcoxon rank-sum test and Mann-Whitney U-test were performed using the wilcoxon and mannwhitneyu functions from scipy.stats in Python. The ggplot2 package in R (r-project.org) was utilized to generate stacked bar plots for model success rate assessment.

### Complex model accuracy assessment

We assessed model accuracy using DockQ^44^, which was downloaded from GitHub (https://github.com/bjornwallner/DockQ). Peptide-HLAII complexes model accuracy was computed by DockQ using the experimentally determined complex structures obtained from the RCSB PDB database. DockQ calculates the fraction of native contacts (fnat), interface backbone RMSD (I- RMSD), ligand RMSD (L-RMSD), DockQ score, as well as the Critical Assessment of PRediction of Interactions (CAPRI) accuracy level, which classifies the model into one of four different accuracy classes: incorrect, acceptable, medium, and high, based on the model’s similarity to the native complex structure.

### Peptide-HLA cross-linking

All cross-linking reaction mixtures were analyzed in two replicates for each tested enzymatic condition (trypsin and trypsin-chymotrypsin digestion). A model CLIP/HLA-DRB1*01:01 complex (4 µM final concentration) was let conditioning at room temperature for 15 min before triggering the cross-linking reactions. PBS (pH ∼7.4) was used as cross-linking reaction buffer. Two different cross- linkers were applied. DSBU cross-linker was dissolved in neat dimethyl sulfoxide (DMSO) and added to the sample to a final concentration of 0.4 mM. The solution was incubated at room temperature for 60 minutes and quenched with ammonium bicarbonate (ABC, 40 mM final concentration) for 15 minutes. DMTMM cross-linker was dissolved in MilliQ water and added to the sample to a final concentration of 361 mM. The solution was then incubated at 37°C for 120 minutes and quenched with ammonium bicarbonate (ABC, 40 mM final concentration) for 15 minutes.

### Enzymatic in-solution digestion

Quenched cross-linked samples (2µg of total protein) were then diluted in Rapigest SF™ (Waters, Milford, USA) 0.1% p/v in 50 mM ammonium bicarbonate (ABC) and reduced with 50 mM dithiothreitol (DTT) in 50 mM ABC for 30 minutes at 60°C. Tubes were cooled down to room temperature before the alkylation step using 200 mM Iodoacetamide (IAA) solution in 50 mM ABC for 20 minutes in the dark at room temperature. Samples were then digested using trypsin digestion or combined trypsin-chymotrypsin digestion. Trypsin digestion (protein to enzyme ratio 20:1) using freshly prepared trypsin (Sigma Aldrich, St. Louis, USA) was performed at 37 °C overnight. For combined digestion, overnight trypsin digestion was followed by chymotrypsin (Thermo Fisher Scientific, Waltham, USA) digestion (protein to enzyme ratio 20:1) for 4 hours at 37°C in a Thermomixer. The obtained peptide mixture was evaporated in a Genevac evaporator, and the final dried samples were resuspended in an aqueous solution containing 2% (v/v) ACN 0.1% (v/v) TFA and subjected to LC-MS/MS analysis.

### Cross-links data analysis

Identification of cross-links was performed with MeroX 2.0.1.4. All MeroX settings are summarized in Supporting Information (SI). MS data have been deposited to the ProteomeXchange Consortium via the PRIDE partner repository with the project accession PXD055226 (Username, Password: mr59EsfXOpTo).

### LC-MS/MS analysis

Cross-linked samples were loaded onto an Acclaim PepMap C18 trap column (300 μm i.d. × 5 mm, Thermo Fisher Scientific, 160454) and separated on a nanocapillary liquid chromatography system (Dionex UltiMate 3000 RSLC, Thermo Fisher Scientific, CA, USA) using a C18 reversed-phase column set at 35°C (75 μm i.d.× 150 mm, Pep Map™ RSLC C18, 2μm, 100Å, Thermo Fisher Scientific, ES904) connected to a Q-Exactive Plus Orbitrap mass spectrometer (Thermo Fisher Scientific) via nanoelectrospray ionization (EasySpray source, Thermo Fisher Scientific). Details are in SI.

## Results

### Cross-link dataset plays a crucial role in the AlphaLink2 model accuracy for all investigated peptide-HLA complexes

To perform a detailed assessment of the impact of the cross-link dataset on the prediction accuracy of AlphaLink2, we selected a subset of 35 different peptide-HLA complexes from the whole dataset. We then defined 4 different simulated cross-link dataset categories: Full represents a cross-link dataset using three different anchor points (peptide N-terminus, peptide C-terminus and a potential reactive amino acid in the core of the peptide binding motif). 2 (Nterm + Cterm) cross-link dataset makes use of two cross-link restraints (one on peptide N-terminus and the other on peptide C-terminus). Nterm and Cterm cross-link datasets involved one cross-link restraint on N- and C- peptide termini respectively. For each cross-link category, top 3 models were predicted using AlphaLink2 and ranked according to their respective model confidence. Interestingly we found a statistical difference in the Wilcoxon signed-rank test when comparing the median values from both L-RMSD and I-RMSD (Figures 1A, B).

**Figure 1.**
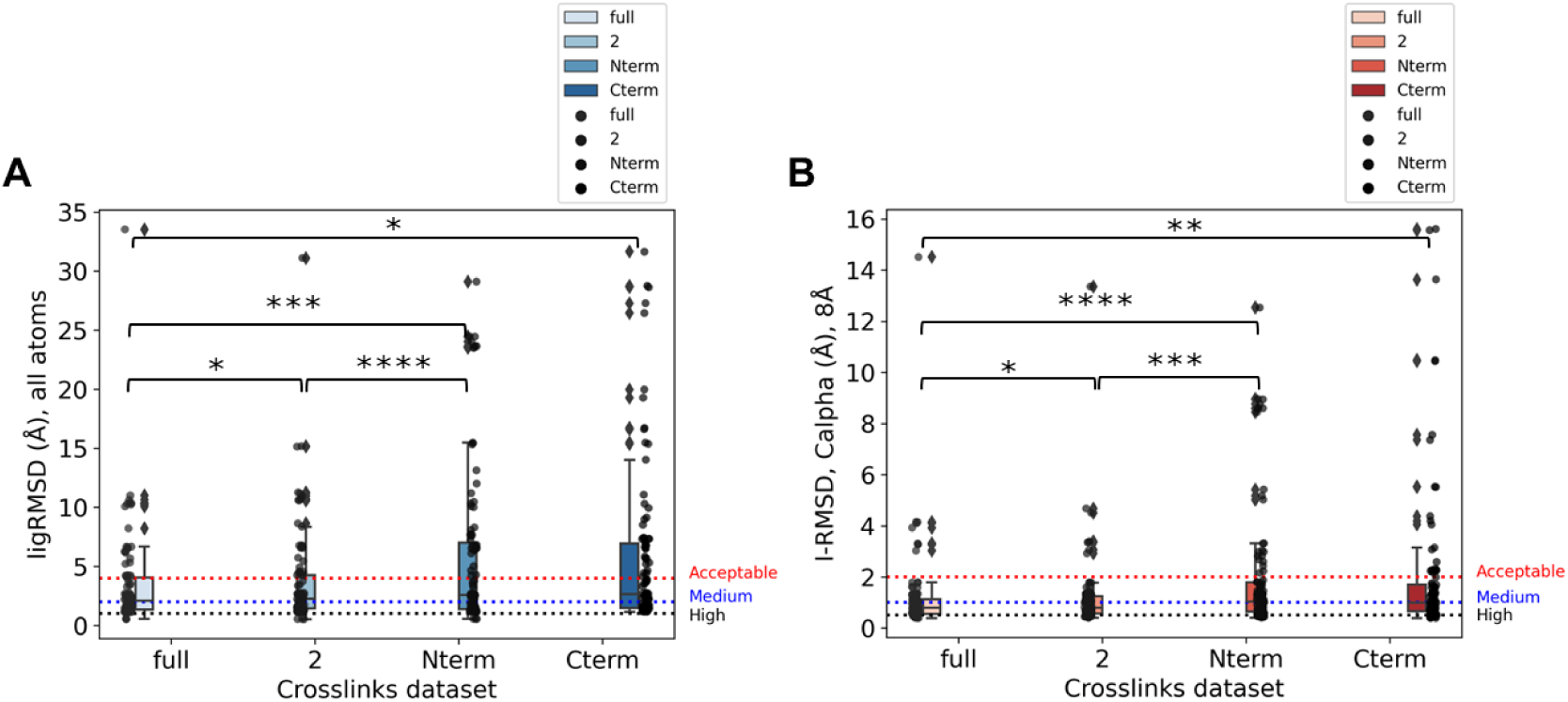
Impact of cross-link dataset on the L-RMSD and I-RMSD of investigated peptide-HLA complexes subset. (A) L-RMSD boxplot distribution for top3 models according to different evaluated cross-link datasets (full, 2, Nterm and Cterm) in AlphaLink2. **(B)** I-RMSD boxplot distribution for top3 models according to different evaluated cross-link datasets (full, 2, Nterm and Cterm) in AlphaLink2. Statistical significance values (Wilcoxon rank-sum test) were calculated for L-RMSD and I-RMSD between different datasets (*p ≤ 0.05; **p≤ 0.01; ***p ≤ 0.001; ****p≤ 0.0001). As horizontal asymptotes are shown the different thresholds for L-RSMD and I-RMSD stated by CAPRI.

Firstly, distributions of both L-RMSD and I-RMSD values of all cross-link datasets followed a nonnormal distribution as assessed by Shapiro-Wilk normality test and Q-Q plot analysis (p-value <0.05, Figures S1, S2). This was also confirmed when plotting the different value distributions in the Culley and Frey graph, showing a typical beta distribution for our observed values (Figures S1, S2). Then, different categories were compared for their median differences using Wilcoxon test or Mann-Whitney U test. Briefly for all-atoms L-RMSD, full and 2 cross-link datasets outperformed Nterm and Cterm dataset (full vs Nterm p-value <0.001, full vs Cterm p-value <0.05, 2 vs Nterm p-value <0.0001, no statistical difference was observed between 2 and Cterm datasets), highlighting the weight of cross-links information in modeling peptide backbone and sidechains (for all peptide no advantage is gained from MSA step of the prediction). Therefore, this was also reflected in the I- RMSD calculation for the different evaluated datasets. More precisely for I-RMSD, full and 2 cross- link datasets still outperformed N and Cterm dataset (full vs Nterm p-value <0.0001, full vs Cterm p-value <0.01, 2 vs Nterm p-value <0.001, no statistical difference was observed between 2 and Cterm datasets).

### Peptide length impact on AlphaLink2/AlphaFold2 models accuracy

To identify potential factors associated with modeling outcome, we analyzed the properties of the native peptide–HLA complexes in relation to predictive modeling success. It is well known from the literature that the frequency of different peptide lengths that can be bound to HLAII follows a normal distribution, where 15 and 16mers outweigh the other peptide length. This is also experimentally confirmed for eluted HLAII ligands in mass spectrometry (MAPPs)^29^. In Figures 2A-H, a trade-off between AlphaLink2 model accuracy (for L-RMSD and I-RMSD) and peptide length is shown, where the model success tended to be lower when peptide length was longer than 16 amino acids. This was also confirmed in AlphaFold2 benchmarking by different peptide lengths (Figures 2I, L). Here, the model accuracy tended to be worse when peptide length was longer than 15 amino acids.

**Figure 2.**
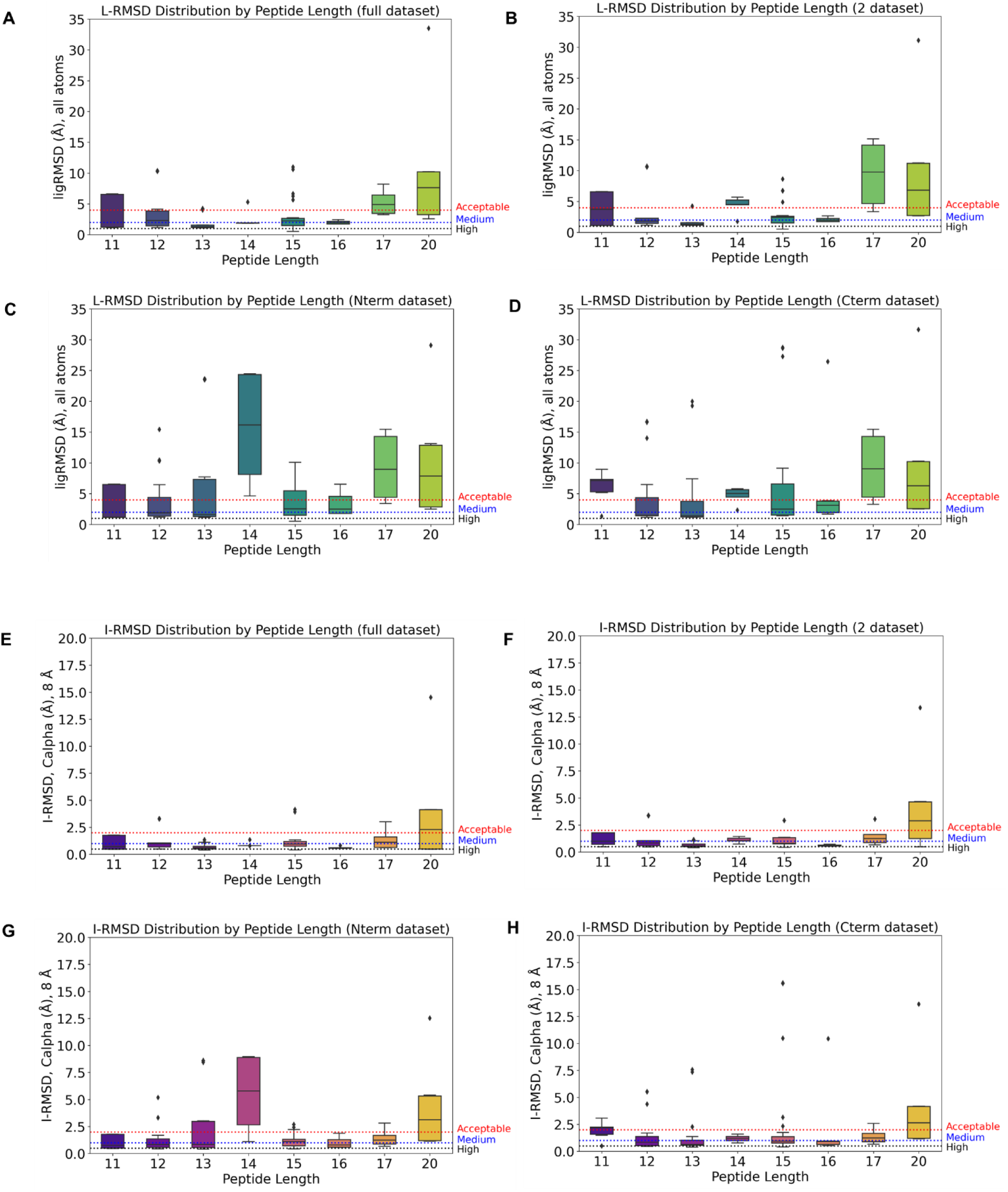
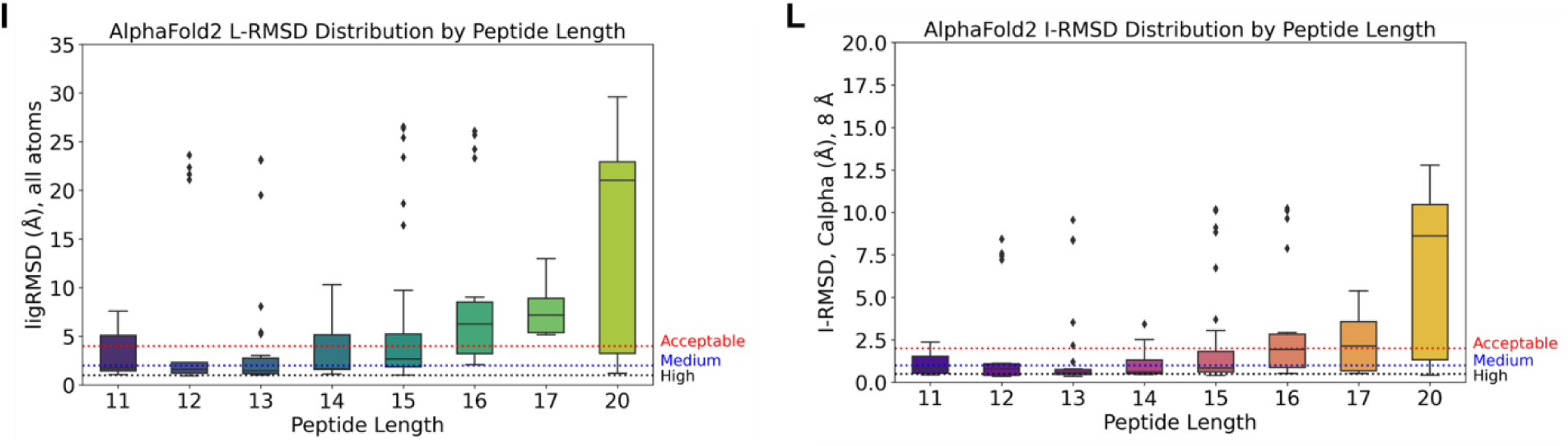
Impact of peptide length on the L-RMSD and I-RMSD of investigated peptide-HLA complexes subset. (A) AlphaLink2 L-RMSD boxplot distribution for top3 models according to different peptide lengths of investigated peptide-HLA dataset for full cross-link dataset. **(B)** AlphaLink2 L-RMSD boxplot distribution for top3 models according to different peptide lengths of investigated peptide-HLA dataset for 2 cross-link dataset (Nterm + Cterm). **(C)** AlphaLink2 L-RMSD boxplot distribution for top3 models according to different peptide lengths of investigated peptide-HLA dataset for Nterm cross-link dataset. **(D)** AlphaLink2 L-RMSD boxplot distribution for top3 models according to different peptide lengths of investigated peptide-HLA dataset for Cterm cross- link dataset. **(E)** AlphaLink2 I-RMSD boxplot distribution for top3 models according to different peptide lengths of investigated peptide-HLA dataset for full cross-link dataset. **(F)** AlphaLink2 I- RMSD boxplot distribution for top3 models according to different peptide lengths of investigated peptide-HLA dataset for 2 cross-link dataset (Nterm + Cterm). **(G)** AlphaLink2 I-RMSD boxplot distribution for top3 models according to different peptide lengths of investigated peptide-HLA dataset for Nterm cross-link dataset. **(H)** AlphaLink2 I-RMSD boxplot distribution for top3 models according to different peptide lengths of investigated peptide-HLA dataset for Cterm cross-link dataset. As horizontal asymptotes are shown the different thresholds for L-RSMD and I-RMSD stated by CAPRI. **(I)** AlphaFold2 L-RMSD boxplot distribution for top3 models according to different peptide lengths of the whole investigated peptide-HLA dataset. As horizontal asymptotes are shown the different thresholds for L-RSMD stated by CAPRI. **(L)** AlphaFold2 I-RMSD boxplot distribution for top3 models according to different peptide lengths of the whole investigated peptide-HLA dataset. As horizontal asymptotes are shown the different thresholds for I-RMSD stated by CAPRI.

### AlphaLink2 vs AlphaFold2 model accuracy for different HLA alleles

Next, we compared AlphaLink2 benchmarking results (full cross-link dataset) with those obtained from AlphaFold2 (in multimer mode). This is important for understanding the benefits of introducing simulated/experimental cross-link restraints to bias peptide-HLA model generation, as MSA information for the peptide is sparse or absent. For fairness of comparison with the full dataset AlphaLink2 pipeline’s results, only AlphaFold2 top 3 predictions were used (Figure 3A).

**Figure 3.**
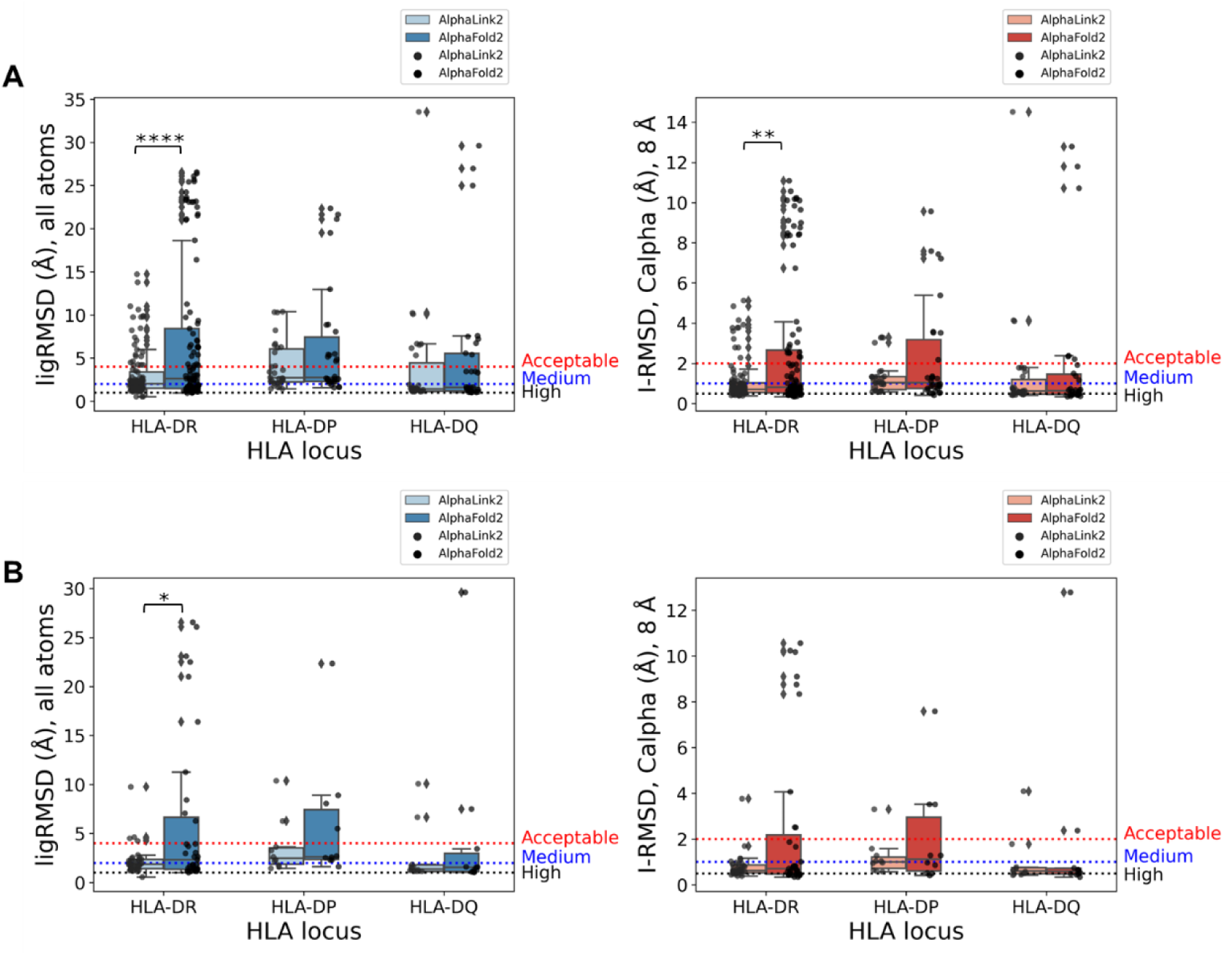
Comparison of AlphaLink2 and AlphaFold2 benchmarking results for peptide-HLA modeling for the three HLAII loci. (A) L-RMSD and I-RMSD comparison for different HLAII allele loci for top3 predicted models (n=105 for peptide/HLA-DR complexes, n=30 for peptide/HLA-DP and DQ complexes, for each modeling pipeline). **(B)** L-RMSD and I-RMSD comparison for different HLAII allele loci for top1 predicted models (best predicted models in terms of model confidence). Statistical significance values (Wilcoxon rank-sum test) were calculated for L-RMSD and I-RMSD between the two computational pipelines for each HLA allele locus (*p ≤ 0.05; **p≤ 0.01; ***p ≤ 0.001; ****p≤ 0.0001).

As shown in Figure 3A, AlphaLink2 performed quite similarly to AlphaFold2 when comparing L- RMSD and I-RMSD for top3 models on HLA-DP (n=30) and DQ alleles (n=30). However, AlphaLink2 performance was better than AlphaFold2 on HLA-DR alleles (n=105). In fact, a statistical difference was found in L-RMSD (p-value <0.0001) and I-RMSD (p-value <0.01) distributions. Notably, a statistical difference remained when comparing median L-RMSD values for best predicted models (top1, p-value < 0.05). In particular, the advantage of introducing simulated cross-link restraints was extremely helpful when correctly predicting noncanonical peptide binding modes. Besides, most commonly for HLA-DP molecules but not limited to them, peptide can bind with an inverse (C-terminus towards P1 pocket, near 310 helix) or flipped conformational binding mode. As shown in Figures 4A-C, three representative case studies predictions were shown in comparison to crystal native peptide binding poses with inverted/flipped peptide binding motifs. Here, introducing physical restraints helped AlphaLink2 to predict the correct peptide binding pose when it is bound in a non-canonical mode.

**Figure 4.**
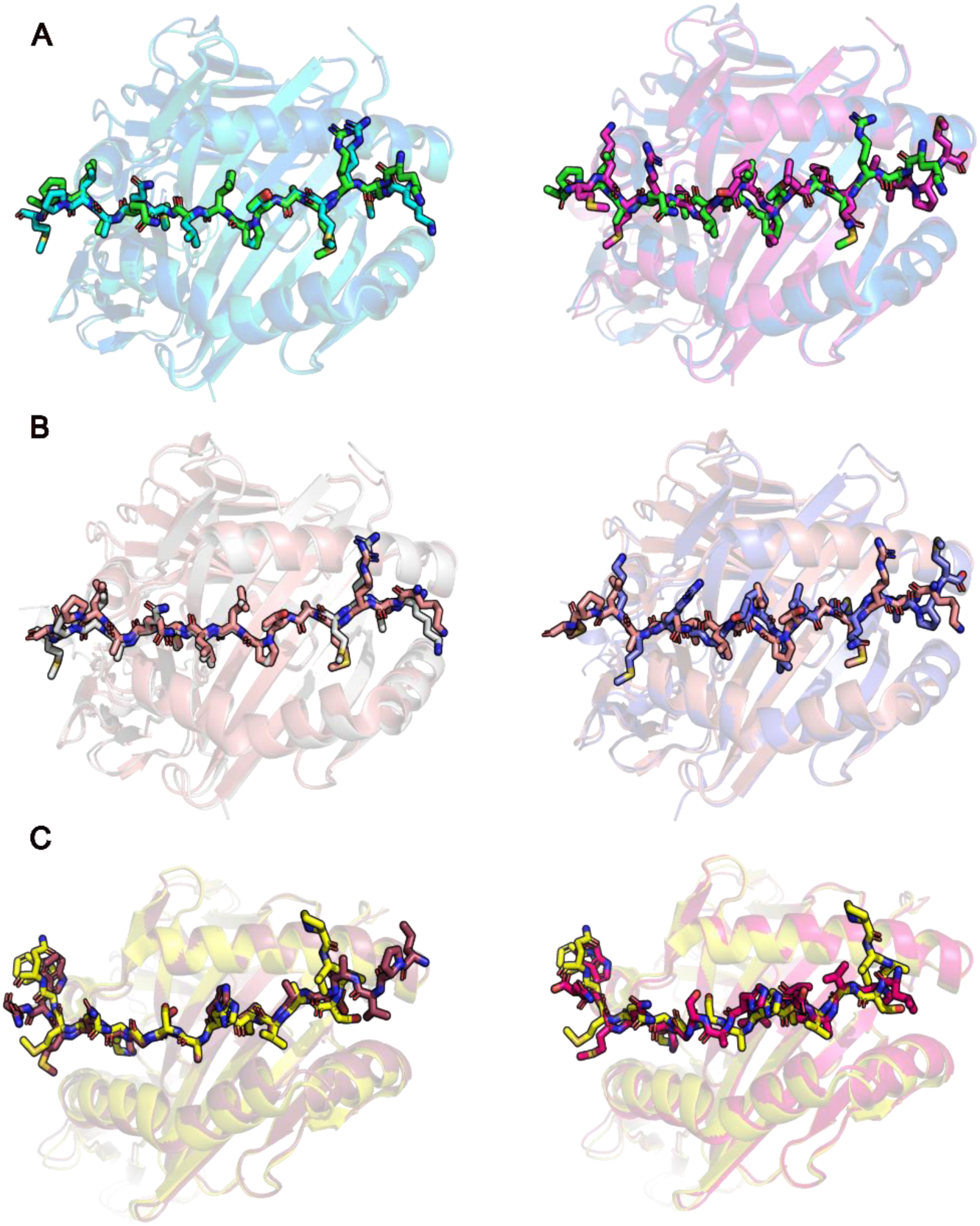
AlphaLink2/AlphaFold2 model accuracy compared to native structures for HLAII ligands with inverted/flipped binding motifs. (A) Example of a near-native prediction by AlphaLink2/AlphaFold2 compared to experimentally determined structure (PDB ID: 4AEN). This AlphaLink2 model has high CAPRI accuracy (DockQ score = 0.90, L-RMSD = 1.76 Å, I-RMSD = 0.55 Å). AlphaFold2 prediction has incorrect CAPRI accuracy (DockQ score = 0.10, L-RMSD = 26.08 Å, I-RMSD = 10.24 Å). AlphaLink2 (left, cyan cartoons) and AlphaFold2 (right, magenta cartoons) models’ superimposition with native structure (green cartoons) for CLIP peptide in reversed peptide orientation with HLA-DR1. **(B)** Example of a near-native prediction by AlphaLink2/AlphaFold2 compared to experimentally determined structure (PDB ID: 3PGC). This AlphaLink2 model has high CAPRI accuracy (DockQ score = 0.91, L-RMSD = 1.49 Å, I-RMSD = 0.75 Å). AlphaFold2 prediction has incorrect CAPRI accuracy (DockQ score = 0.10, L-RMSD = 26.55 Å, I-RMSD = 10.18 Å). AlphaLink2 (left, grey cartoons) and AlphaFold2 (right, purple cartoons) models’ superimposition with native structure (orange cartoons) for CLIP peptide in flipped conformation with HLA-DR1. **(C)** Example of a near-native prediction by AlphaLink2/AlphaFold2 compared to experimentally determined structure (PDB ID: 7T6I). This AlphaLink2 model has medium CAPRI accuracy (DockQ score = 0.78, L-RMSD = 3.63 Å, I-RMSD = 0.64 Å). AlphaFold2 prediction has medium CAPRI accuracy (DockQ score = 0.54, L-RMSD = 5.50 Å, I-RMSD = 1.1 Å). AlphaLink2 (left, brown cartoons) and AlphaFold2 (right, pink cartoons) models’ superimposition with native structure (yellow cartoons) for pp65 peptide in reverse peptide orientation in complex with HLA-DP1.

### AlphaLink2/AlphaFold2 prediction accuracy on single HLA alleles

To better understand the factors that can contribute to the success rate of the AlphaLink2/AlphaFold2 peptide-HLA modeling, we investigated the performance of both pipelines on single HLA alleles loci and compared them (Figures 5A, B). We observed higher success in modeling peptide/HLA-DR and DQ complexes compared to HLA-DP for AlphaLink2 top3 predictions, in terms of L-RMSD (p-value <0.01 for HLA-DR vs HLA-DP and <0.001 for HLA-DQ vs HLA-DP) and I-RMSD (p-value < 0.01 for both HLA-DR and HLA-DQ). This was also confirmed from AlphaFold2 benchmarking results, where also peptide/HLA-DP complexes were predicted with minor accuracy compared with HLA-DQ (p-value < 0.05). This could be explained by the fact that there might be fewer peptide/HLA-DP published crystal structures in the training set than HLA-DR and DQ. Another explanation for the lower accuracy of peptide/HLA-DP predictions could be explained by more common “inverted” binding motifs of these HLA molecules compared to the classical peptide binding mode in HLA-DR and DQ molecules.

**Figure 5.**
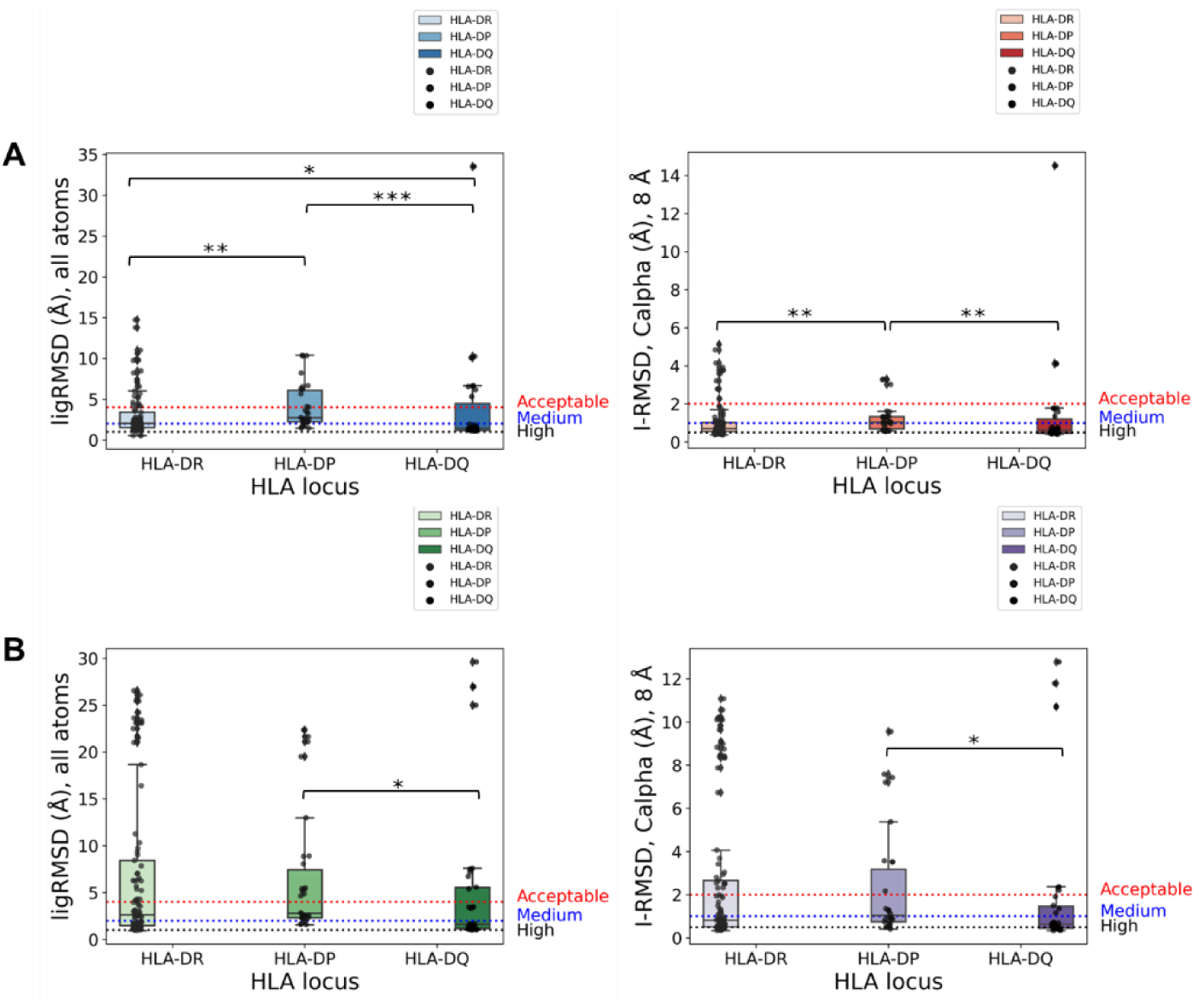
AlphaLink2/AlphaFold2 performances on single HLA alleles. (A) AlphaLink2 performance comparison in terms of L-RMSD and I-RMSD (top3 predictions) for the investigated HLA loci (n=35 unique HLA-DR alleles, n=10 unique HLA-DP and DQ alleles). **(B)** AlphaFold2 performance comparison in terms of L-RMSD and I-RMSD (top3 predictions) for the investigated HLA loci (n=35 unique HLA-DR alleles, n=10 unique HLA-DP and DQ alleles). Statistical significance values (Mann-Whitney U test) were calculated for L-RMSD and I-RMSD between the different HLA allele locus categories for each computational pipeline (*p ≤ 0.05; **p≤ 0.01; ***p ≤ 0.001; ****p≤ 0.0001).

### Model confidence scores comparison to infer near-native predictions

Since the good outcome of model accuracy scores produced by AlphaFold2^40,45^ we also wanted to evaluate the ability of those scores, or adjustments of these, to discriminate between accurate versus incorrect peptide-HLA predictions. We assessed AlphaFold2/AlphaLink2 model confidence score, which is a linear combination of pTM and ipTM^38,40^ scores, as well as interface pLDDT (I-pLDDT), which is based on residue-level confidence scores for peptide-HLA interface residues (8 Å distance cutoff), as used in previous studies^45^, for discriminating correct peptide-HLA models according to different CAPRI DockQ categories. Model confidence values did not enable to discriminate high-medium accuracy models compared to inaccurate models properly (AlphaFold2 model confidence range spanned from 0.86 to 0.94, AlphaLink2 model confidence from 0.86 to 0.93), even though high-quality models tended to have higher model confidence values compared to incorrect models (Figures S3, S4). Interestingly, I-pLDDT showed a similar correlation to DockQ scores, but it allowed a better discrimination between accurate and incorrect models (AlphaFold2 I-pLDDT range spanned from 78.2 to 98.1, AlphaLink2 I-pLDDT from 77.3 to 98.0, Figures S3, S4). Both of these values are biased by the largely dominant amino acid sequence of HLA protein, which is always well-predicted by AlphaLink2 and AlphaFold2. We also tested the individual components of the model confidence scores (pTM and ipTM) (Figures S3, S4), which did not yield improved correlations with DockQ scores and had the same dynamic range interval of model confidence. Furthermore, we also assessed the correlation between peptide LDDT (pepLDDT) and DockQ accuracy scores and L-RMSD (Figure S5). Surprisingly, we found an interesting correlation between pepLDDT and DockQ/L-RMSD, and it also provided outstanding discrimination between incorrect versus medium or higher accuracy models for AlphaLink2 and AlphaFold2 (Figures 6A, B). This suggests that pepLDDT score can potentially discriminate corrected peptide binding poses from unplausible binding conformations.

**Figure 6.**
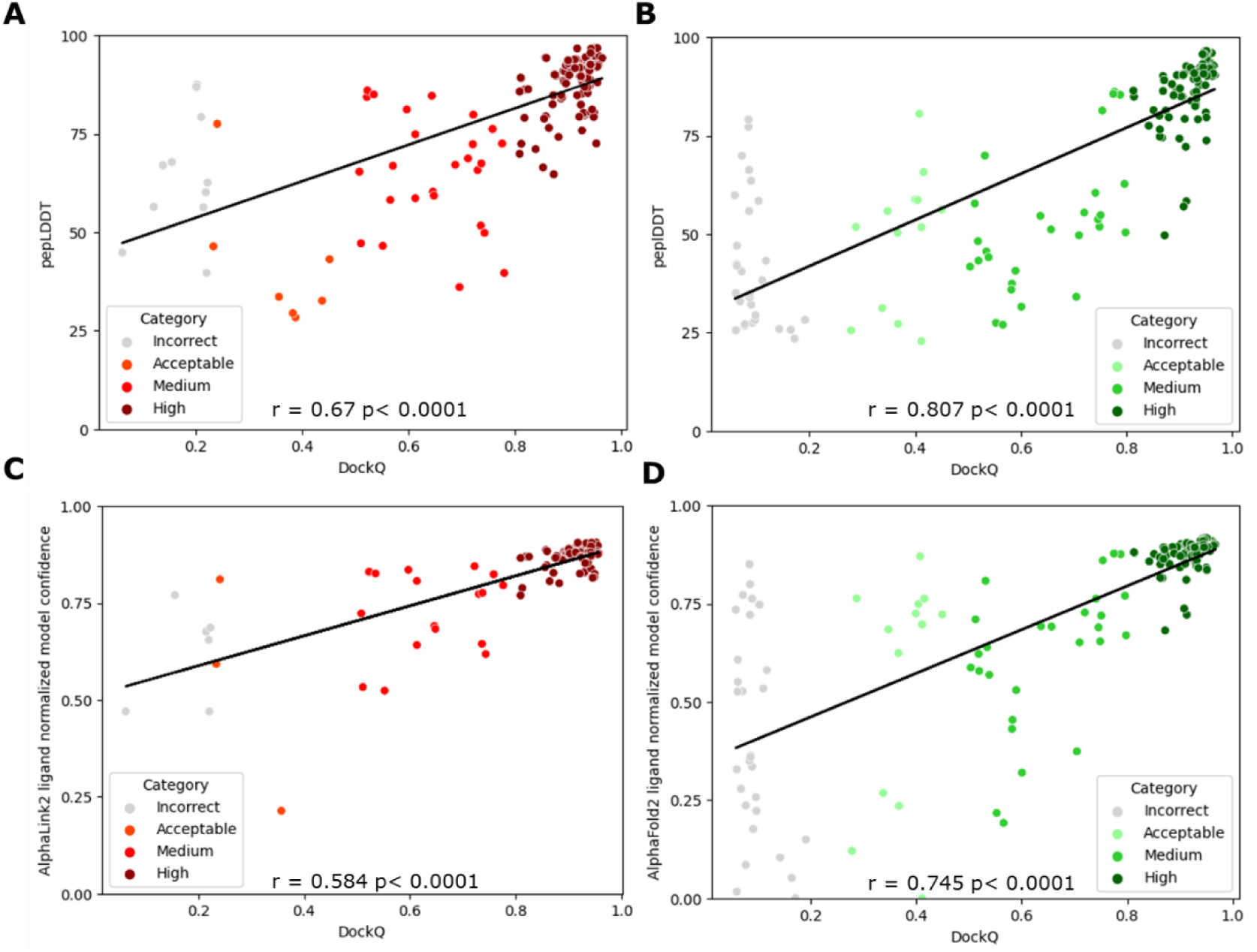
AlphaLink2/AlphaFold2 model predictor scores and DockQ model accuracy correlations. Scatter plots correlation of pepLDDT for AlphaLink2 **(A)** and AlphaFold2 **(B)** score with model accuracy, assessed by DockQ score. Scatter plots correlation of ligand normalized model confidence for AlphaLink2 **(C)** and AlphaFold2 **(D)** score with model accuracy, assessed by DockQ accuracy score. In the scatter plots, all models representing 165 complexes are depicted as data points, with their colors indicating the model quality according to CAPRI criteria. The black line represents the linear regression, and in the lower center of the scatter plots Pearson’s correlation coefficients and correlation p-values are displayed for each model predictor.

To weight the peptide backbone and side-chains conformation into the model confidence, a ligand normalization score (LNS) was used to derive the ligand normalized model confidence from original model confidence values (Equation 1-2, Figures 6C, D). Remarkably, we found a similar correlation of ligand normalized model confidence with DockQ score and L-RMSD (Figures S5, S6) compared to pepLDDT for AlphaLink2/AlphaFold2. Moreover, it also provided better discrimination between incorrect versus medium or higher accuracy models for both AlphaLink2 and AlphaFold2 compared to the original model confidence scores (Figures 6C, D). The Kernel density distribution of pepLDDT and ligand normalized model confidence scores for AlphaLink2 and AlphaFold2 is shown in Figures 7A-D.

**Figure 7.**
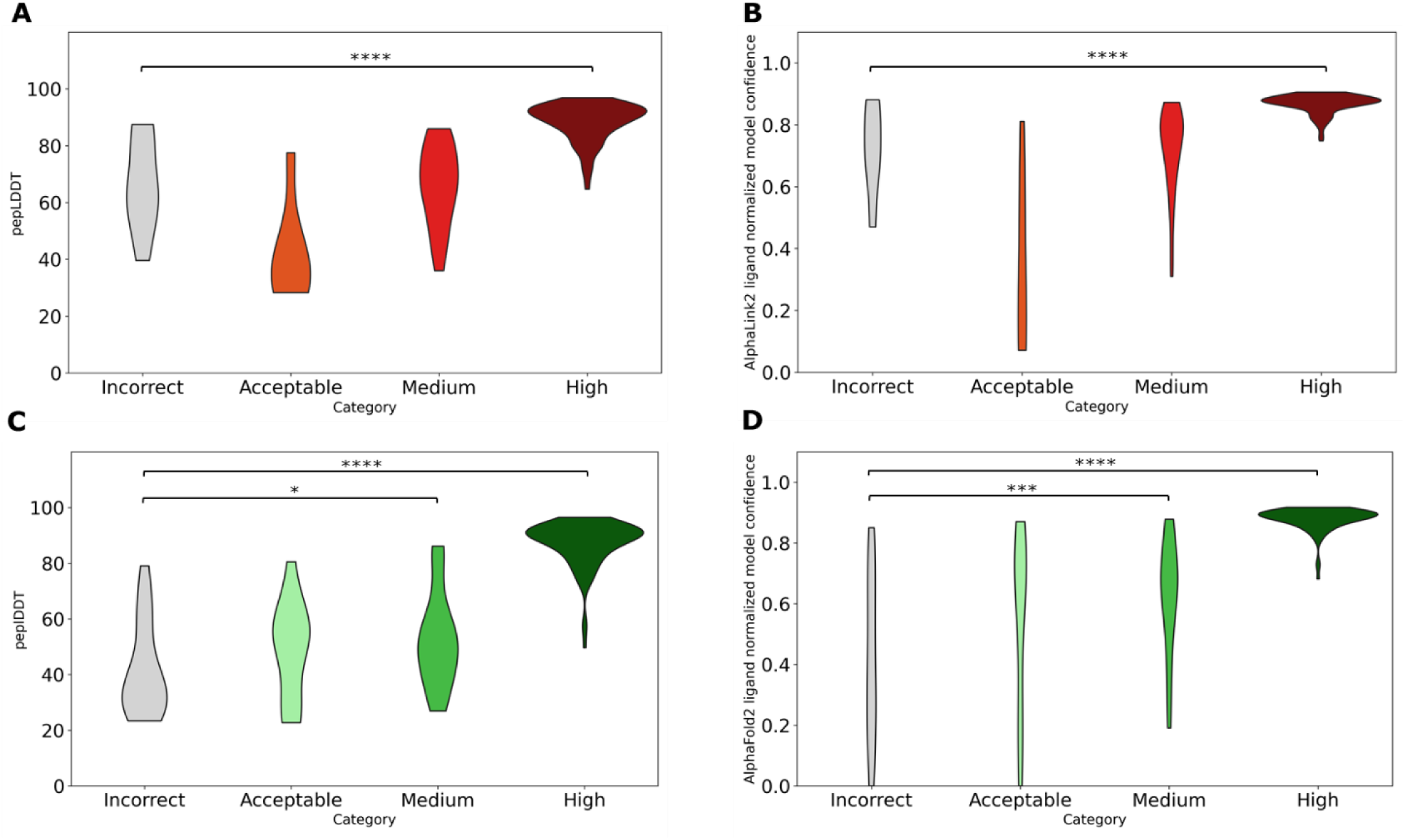
AlphaLink2/AlphaFold2 model predictor scores Kernel distribution and DockQ model accuracy. Kernel distribution of **(A, C)** pepLDDT and **(B, D)** ligand normalized model confidence, grouped by the CAPRI criteria for AlphaLink2 (A, B) and AlphaFold2 (C, D) top3 predictions. Statistical significance values (Mann-Whitney U test) were calculated between model scores for sets of predictions with incorrect versus medium and incorrect versus high CAPRI accuracy, as noted at top (* p ≤ 0.05; **p≤ 0.01; ***p ≤ 0.001; **** p≤ 0.0001).

In Figure 7A, the distribution of peptide LDDT scores according to different DockQ categories for AlphaLink2 is shown. A significant difference was found between the Incorrect category and High category (p-value <0.0001). In Figure 7C, pepLDDT scores distribution for AlphaFold2 is reported. This plot also showed significant difference between the Incorrect category and higher categories (p-value < 0.0001 for High accuracy and <0.05 for Medium accuracy models). In Figure 7B, AlphaLink2 ligand normalized model confidence score distribution according to different accuracy categories is shown. Notably, ligand normalized confidence scores still increase from Incorrect to High category, with a significant difference (p-value < 0.0001). The same conclusions can be drawn for AlphaFold2 ligand normalized model confidence distribution along the different DockQ categories (p-value < 0.0001 for High accuracy and <0.001 for Medium accuracy models, Figure 7D).

Overall, this demonstrated that both AlphaLink2 and AlphaFold2 model predictor scores perform better as the model quality improves from Incorrect to High. The significant difference in scores emphasized a good discrimination between higher accuracy and low accuracy predictions for the HLA complexes tested in this study. In fact, they also provided outstanding discrimination between incorrect versus medium or higher accuracy models based on receiver operating characteristic (ROC) area under the curve (AUC) metrics (AlphaFold2 ligand normalized model confidence/pepLDDT AUC = 0.97, AlphaLink2 ligand normalized model confidence AUC = 0.92, AlphaLink2 pepLDDT AUC = 0.94), which are higher than that of the respective model confidence (AlphaFold2 model confidence AUC=0.95, AlphaLink2 model confidence AUC = 0.87, Figures S7A-D). AlphaFold2’s Acceptable category showed a broader range of scores compared to AlphaLink2, but both pipelines showed a clear trend of increasing accuracy and confidence in better categories (Figures 7A-D). This indicates that these tools effectively distinguish between less and more accurate peptide structure predictions.

### Evaluating a cross-linker combination to characterize a model CLIP peptide-HLA DRB1*01:01 complex

To evaluate the AlphaLink2 performance with experimentally-derived cross-links, we assessed a combination of two different cross-linkers on a model CLIP peptide-HLA molecule. Specifically, DSBU and DMTMM were used to perform cross-linking reactions. We identified 16 distinct intermolecular cross-links between the peptide and the HLA molecule (Table S2, Figures 8A, C), from which a representative fragment ion mass spectrum is shown in Figure 8B. These cross-links were identified between the peptide N-terminus and a threonine in the peptide core, that interact with different residues of the variable regions of α1 and β1 HLA chains (Table S2). AlphaLink2 generated top1 model was superimposed with an experimentally-determined CLIP-HLA crystal structure, showing an all-atoms RMSD of 1.99 Å (Figure 8D, Figure S8A), that was better compared to AlphaFold2 top1 model (all-atoms RMSD = 2.33 Å, Figure S8B). The AlphaLink2 model showed a peptide L-RMSD (all-atoms) of 1.30 Å compared to native structure (AlphaFold2 top1 model L- RMSD = 1.69 Å, Figure S8B).

**Figure 8.**
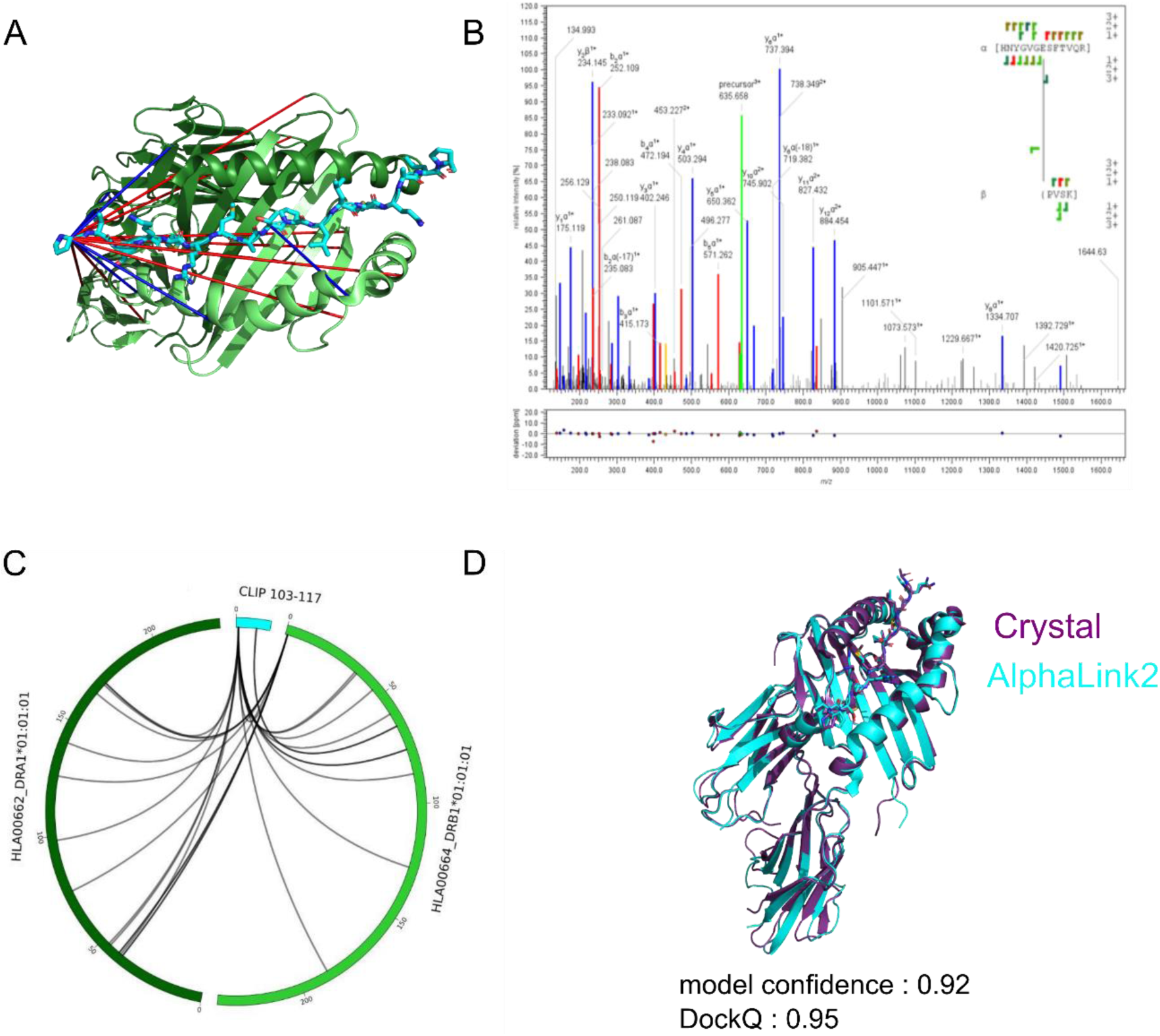
**Cross-linking mass spectrometry and integrative modeling on CLIP-HLA DRB1*01:01 results summary**. **A)** CLIP (103-117)-HLA DRB1*01:01 intermolecular protein cross-links are mapped on a crystal structure of bound complex (PDB ID: 3PDO). 40% of these cross-links were satisfied (Cα-Cα distance ≤ 25.0 Å. **B)** Representative fragment ion spectrum of an intermolecular cross-link between E87 of β1 chain (HNYGVG<u>E</u>SFTVQR, peptide α) and CLIP peptide N-terminus (<u>P</u>VSK, peptide β) DMTMM inter-protein cross-link. Y- and b-type ions series for the α and β peptides are presented in blue and red, respectively. Precursor ion is showed in green. **C)** Circular plot of identified CLIP-HLA inter-protein cross-links using the combination of two crosslinkers and enzymes (see Table S2 for details). **D)** Superimposition between crystal structure (purple cartoon) and AlphaLink2 top1 model (cyan cartoon) using experimentally-obtained cross-links from the tested cross-linker combination. This AlphaLink2 model has high CAPRI accuracy (DockQ score = 0.95, all-atoms RMSD = 1.99 Å, L-RMSD = 1.30 Å, I-RMSD = 0.51 Å). Results summary of inter/intramolecular protein cross-links for each cross-linker is shown in Figure S9 of SI.

### AlphaLink2/AlphaFold2 peptide-HLAII modeling accuracy and success rate

The accuracy of peptide-HLAII complex predictions was evaluated using Critical Assessment of Predicted Interactions (CAPRI) criteria, which classify predictions as incorrect, acceptable, medium, or high based on a combination of interface root-mean-square distance (I-RMSD), ligand root-mean-square distance (L-RMSD), and the fraction of native interface residue contacts (fnat), by comparison with the experimentally-determined complex structure. AlphaLink2 generated medium or higher accuracy models, which we refer to as near-native predictions, as top1-ranked predictions for 94% of the cases, and high accuracy models were generated for 82% of the test cases (Figure 9). AlphaFold2 generated acceptable or higher accuracy models as top1-ranked predictions for 84% of the 55 test cases for which models were generated (Figure 9). Medium or higher accuracy models, which we refer to as near-native predictions, were generated as top-ranked predictions for 75% of the cases, and high accuracy models were generated for 65% of the test cases.

**Figure 9.**
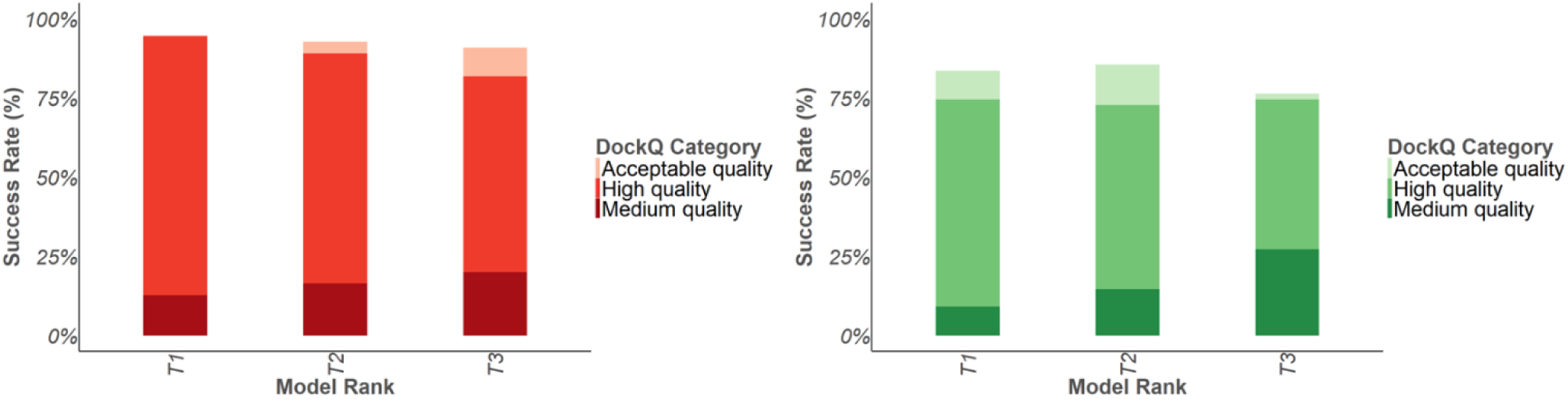
**Peptide-HLAII modeling accuracy of AlphaLink2/AlphaFold2**. **(Left)** Benchmarking of AlphaLink2 was performed on 55 peptide-HLAII complexes. **(Right)** Benchmarking of AlphaFold2 was performed on the same set of peptide-HLAII complexes. For each complex, 3 predictions were generated and ranked by AlphaLink2/AlphaFold2 model confidence score. Peptide-HLAII complex predictions were evaluated for complex modeling accuracy using CAPRI criteria for high, medium, and acceptable accuracy. The success rate was calculated based on the percentage of cases that had at least one model among their top N ranked predictions that met a specified level of CAPRI accuracy. Bars are colored by CAPRI accuracy level.

### Conclusions and future perspectives

Using a set of 55 nonredundant peptide-HLAII complexes, we benchmarked and evaluated the ability of AlphaFold2 and AlphaLink2 to model peptide-HLAII complexes. In this set, we observed a good success rate in predicting peptide-HLAII structures by AlphaFold2 (ColabFold version 1.5.5) and AlphaLink2 (Colab Notebook). AlphaLink2 success rate in top1 model was about 94% (with 94% near-native solutions), and for AlphaFold2 was 84% (with 75% near-native solutions). AlphaLink2 high accuracy models were generated for 82% of the test cases against 65% of AlphaFold2. This highlights the importance of introducing cross-link restraints to bias the model generation and improve the peptide conformational sampling. Our benchmarking also showed that AlphaLink2 Colab Notebook exhibited improved success in predicting peptide-HLAII structures versus AlphaFold2 Colab version (v.1.5.5), especially in case of noncanonical peptide binding motifs. We also found that AlphaLink2 and AlphaFold2 are more successful at modeling peptide-HLAII complexes for HLA-DR and DQ loci and have more difficulty in predicting the structure of peptide/HLA-DP complexes. This might be explained by the limited number of peptide/HLA-DP structures compared to the other HLA loci potentially available in the training set. Moreover, there is a higher probability of having noncanonical peptide conformations that could reduce the success rate. This was confirmed in AlphaLink2 and AlphaFold2 benchmarking on top3 and top1 predictions. Importantly, we also found a good correlation between pepLDDT and DockQ accuracy score (Pearson’s correlation coefficient r = 0.67 for AlphaLink2 and r = 0.81 for AlphaFold2) and excellent prediction power in model classification (ROC AUC = 0.94 for AlphaLink2 and AUC = 0.97 for AlphaFold2 for high quality models). Also, adopting a ligand normalized model confidence score, we were able to achieve a better discrimination between incorrect and acceptable quality models compared to model confidence scores. In conjunction with the observed confidence scoring accuracy, these results indicate that researchers may use AlphaLink2 and AlphaFold2 to model this important class of complexes and can complement or assist their experimental structural determination. Moreover, to our knowledge, this is the first application of cross-linking mass spectrometry and integrative modeling in the field of immunopeptidomics. Using a combination of two cross-linkers, we obtained different interprotein cross-links between the peptide and the HLA binding pocket residues, which led to an improvement of the investigated CLIP (103-117)-HLA complex AlphaLink2 prediction accuracy compared to AlphaFold2 predicted model. In the future, these approaches can drive the choice of other *in vitro*/*silico* assays (such as in-solution mutant peptide-HLAII screening binding assays, MAPPs/DC-T cell assays) and be integrated with *in silico* immunogenicity sequence-based predictions, leading to an integrated strategy to screen and reduce the number of potential peptide mutants/variants for developing new vaccines and mitigate immunogenicity risks of exogenous proteins/peptides and assist their deimmunization process.

## Author Contributions

**Andrea Di Ianni:** Conceptualization; methodology; writing – original draft; writing – review and editing; formal analysis; data curation.

**Luca Barbero:** Conceptualization; writing – review and editing.

**Kyra Cowan:** funding acquisition, writing – review and editing.

**Federico Riccardi Sirtori:** funding acquisition, writing – review and editing.

## Notes

The authors declare the following financial interests/personal relationships, which may be considered potential competing interests: KC, LB, and FRS are employees of Merck KGaA.

## ACKNOWLEDGMENT

This research was supported by Merck KGaA Darmstadt, Germany and the University of Turin.

